# Quantification of spatial subclonal interactions enhancing the invasive phenotype of paediatric glioma

**DOI:** 10.1101/2021.07.19.452844

**Authors:** Haider Tari, Ketty Kessler, Nicholas Trahearn, Benjamin Werner, Maria Vinci, Chris Jones, Andrea Sottoriva

## Abstract

Intra-tumour heterogeneity is an intrinsic property of all cancers. In some cases, such variation can be maintained by interactions between tumour subclones with distinct molecular and phenotypic characteristics. In paediatric gliomas, interactions can take the form of enhanced invasive phenotype, a hallmark of these malignancies. However, subclonal interactions are hard to quantify and difficult to distinguish from spatial confounding factors and experimental bias. Here we combine spatial computational modelling of cellular interactions and invasion, with co-evolution experiments of clonally disassembled primary glioma lines derived at autopsy. We design a Bayesian inference framework to quantify spatial subclonal interactions between molecular and phenotypically distinct lineages with different patterns of invasion. We show how this approach could discriminate genuine subclonal interactions where one clone enhanced the invasive phenotype of another, from apparent interactions that were only due to the complex dynamics of subclones growing in space. This study provides a new approach for the identification and quantification of spatial subclonal interactions in cancer.

## Introduction

Intra-tumour heterogeneity represents a significant barrier to successfully developing cancer therapies due to the emergence of treatment resistance^1^. This is particularly true in the case of paediatric glioblastoma and diffuse intrinsic pontine glioma (DIPG), a heterogeneous group of high-grade gliomas^2^ characterised by a highly invasive phenotype. These malignancies invade the brain parenchyma, making resection difficult and leading to poor prognosis^2^.

Intra-tumour heterogeneity is the natural consequence of an evolutionary process driven by random mutation, neutral drift, and non-random, positive and negative selection^4^. However, as paediatric malignancies maintain a remnant of the differentiation programme, cell signalling leading to interactions between lineages of cells or subclones has also been described^5–8^. Evidence of subclonal interactions in other cancer types have been explored^9–11^. However, these interactions remain difficult to quantify, and experimental observations are subject to bias and unaccounted confounding factors, such as spatial constraints and lack of mechanistic models applied to the data to test different alternative hypotheses.

Subclonal interactions can be studied through the lens of evolution and ecology, which seek to understand the dynamics of a particular population within its environment and in relation to others. Whereas the most evident negative interaction between populations is competition for space and resources, leading to Darwinian selection^12^, there are also other forms of interactions such as amensalism, where the negative effect is only experienced by one population^13,14^. Positive interactions instead lead to a population benefitting from the presence of another, and can arise in three varieties^13,14^ (Figure 1a):

- mutualism where each population benefits from the presence of the other
- commensalism where the benefit is only experienced by one population but a cost is not incurred in the opposite direction
- exploitation where a benefit is experienced by one population and a cost to the other.

**Figure 1:**
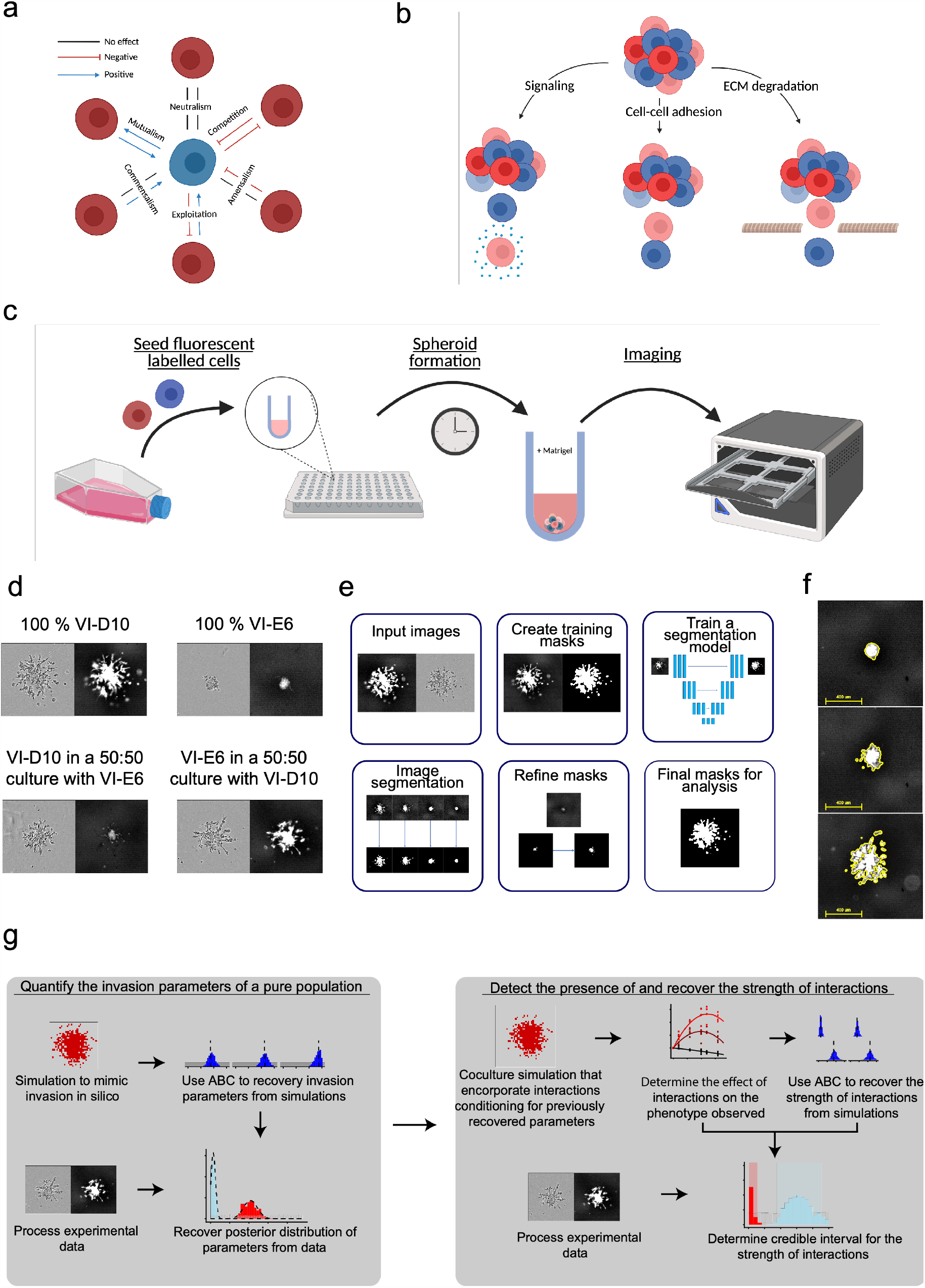
Combining computational modelling with invasion assays to measure cellular interactions. **a**, Illustration of interactions between sub-populations characterised by the affect each population has on each other. **b**, Potential mechanisms through which invasion interactions can occur. **c**, Images of pure populations using phase and GFP channel demonstrates effectiveness labelling whilst 50:50 ratio demonstrates that the phase channel is inadequate in understanding the phenotype of a sub-population. **d**, Invasion assay involves seeding labelled cells to form spheroid, which are embedded in Matrigel. These spheroids are images using the Incucyte S3 live-cell analysis system. **e**, Deep learning image analysis: i) Histogram equalisation and filtering of unfocused images to prepare the input images, ii) Training data set prepared by creating ground truth masks, iii) Training a ResNet-UNet neural network, iv) Segmentation model applied to images, v) Image masks are compared to real images to correct for any false negative pixels (under segmentation). vi) The final binary mask, this is used for further analysis. **f**, Images highlighting the result of image processing, with an outline of the binary mask (yellow) overlaid on the green fluorescence channel. **g**, i) Quantifying the invasion parameters of a pure populations. Using simulations to generate informative measures that are be applied to experimental data to infer parameters. ii) Detecting and quantifying the strength of interactions. Using simulations conditioned with pure population parameters and introducing interactions, measures are developed to understand the effect of interactions on the invasive phenotypes. Applied to data, this generates a credible interval of the true interaction strength.

Exploitation interactions lead to a reduction in the size of the exploited population and can result in its extinction, making these types of interactions rare in cancer where turnover is considerably faster than in species. Mutualism is a two-sided benefit thus requires two species to evolve to occupy complementary niches. Although it could be unlikely to concomitantly evolve this form of adaptation by two independent subclones in the short timescales of a growing malignancy compared to millions of years in natural species, this has been suggested as a potential avenue for the survival of heterogeneous subclones^15^. Commensalism is instead potentially likely to be observed as it requires just one population to provide an interaction that other populations can benefit from. A common example of this could be seen from extracellular signalling through secreted factors, these can be considered as ‘public goods’ where all cells in the tumour benefit from them whilst not all cells produce them^15,16^. Indeed, this has even been seen to confer treatment resistance in pancreatic cancer where TGFa and amphiregulin produced by treatment resistant clones confers resistance to sensitive clones^17^. There have been also some studies that have found interactions driving tumour initiation^18^, metastasis^10^ and cell growth^19^.

Mathematical modelling of cellular population dynamics allows simulating different interaction models and assess whether the observations fit the data. There is vast literature of mathematical models for cancer^20^, however these are not very often applied to data directly with the intent of inferring biology. When this is done the effect is powerful and the resulting conclusions are measurable and translatable to further exploration^21^. Furthermore, although seminal studies have modelled interactions with Evolutionary Game Theory in terms of growth advantage of subclones^22^, these approaches are almost invariably non-spatial, and are not therefore suitable capture the spatial patterns of invasion that we want to study in paediatric gliomas.

In the case of DIPG we are interested in understanding whether there are cellular interactions driving its most prominent phenotype: the diffuse pattern of invasion. There are multiple possible mechanisms via which this interaction could occur; cell-cell adhesion leading to the co-invasion of cells, extracellular matrix degradation allowing for cells lacking this ability to escape and paracrine signalling (Figure 1b)^5^.

In this study leverage on a unique set of primary glioma cell lines derived from patients at autopsy that have been thoroughly characterised at the molecular and phenotypic levels. Importantly, different subclones with distinct molecular features and invasion characteristics have been isolated^5^. We integrate *in vitro* assays of co-evolution between subclones with a spatial computational modelling framework to quantify the presence of spatial subclonal interactions or lack of thereof.

## Results

### Experimental design to recreate invasion in vitro and deep learning based image analysis of invasion

From patient-derived cell lines SU-DIPG-VI, different clones were isolated in 2D and in 3D^5^. High-depth sequencing revealed common mutations between all subclones such as *H3F3A* K27M, *TP53*, PSG5; some mutations were shared such as *NOTCH2, SMARCC*2 and *PHF21*A in VI-D10 and some mutations were private such as *SMARCB1* in VI-E6 only. Their phenotypic characterization in mono-culture revealed that VI-D10 is a fast growing, more invading and migrating subclone *in vitro* compared to VI-E6. Similarly to SU-DIPG-VI, subclones have been isolated in 2D and 3D from HSJD-DIPG-007 patient derived cell line^5^. 007-F10 and 007-F8 share common mutations with the other subclones such as *H3F3A* K27M, *ACVR1, PIK3CA, PPM1D*. Interestingly 007-F10 showed a private mutation R187*coding for a codon stop in the histone lysine transferase gene *KMT5B* detected in 0.3% of the tumour bulk cells. Its isogenic clone 007-F8 is wild-type for *KMT5B*. 007-F10 was characterized by a decrease of H4K20me2, and an increase of invasion/migration *in vitro* over 007-F8.

We grow these clones in a mono-culture and co-culture them at various ratios to assess the differences in the invasive phenotype intrinsic to each clone, as well as the difference in the invasive phenotype between the same subclonal population at different ratios in a co-culture. (Figure 1c).

The experimental design intends to mimic cellular infiltration *in vitro* whilst being able to track individual invading cells. Cell labelling allows for the identification of a specific subclone in a co-culture, where the phase image channel shows all cells, the green channel shows the presence of only the labelled subclone (Figure 1c). To this end, we stably labelled cells with a green fluorescent marker, which we then grew as single spheroids in ultra-low attachment round bottom wells. This allowed for the formation of spheroids which we then embedded in a gelatinous matrix, consisting of structural proteins and growth factors, designed to mimic the extracellular matrix found in the brain. The embedded spheroid was then imaged over 3 days using the Incucyte S3 live-cell analysis system, which produced data in the form of images every 24 hours (Figure 1d).

We used a tile-based deep learning image segmentation algorithm to process the images. These final marks are then compared to the ground truth and corrected to ensure accurate segmentation (Figure 1e). The result of this is to create binary masks, which we can use for image quantification and inference. The masks can be overlaid on the original image as an outline to highlight the segmentation results (Figure 1f).

### Combining computational modelling with invasion assays to measure cellular interactions

Spatial simulations based on cellular automata (CA) allow for the exploration of a wide range of parameters leading to different spatial patterns expected in the data. Statistical inference based on Approximate Bayesian Computation^23^ then allows fitting the model parameters to the data, thus quantifying biological characteristics from imaging of the cell cultures (Figure 1g). The inference results can be verified *in silico* to ensure the correct parameters are recovered from simulated data (see Material and Methods for details).

We start by inferring parameters for a monoclonal population, to then create a null model of invasion in mixed cultures. From this ‘baseline’ model we introduce subclonal interactions to test deviations from the ‘null’. We determine a method of recovering the strength of interactions in silico and apply this method *to in vitro* data to measure the presence and strength of interactions in our experimental system (Figure 1g).

### Simulating invasion

An agent-based cellular automaton (CA) was used to recreate cellular invasion *in silico*. The phenotype of each cell is divided into two categories; growth and movement. These are chosen so as to not overcomplicate models, allowing for this simulation to be applicable to a wide variety of invasion experiments. We use Gillespie’s stochastic simulation algorithm^24^ in order to generate simulations that will have temporal dynamics that are comparable to real data. (Figure 2a).

**Figure 2:**
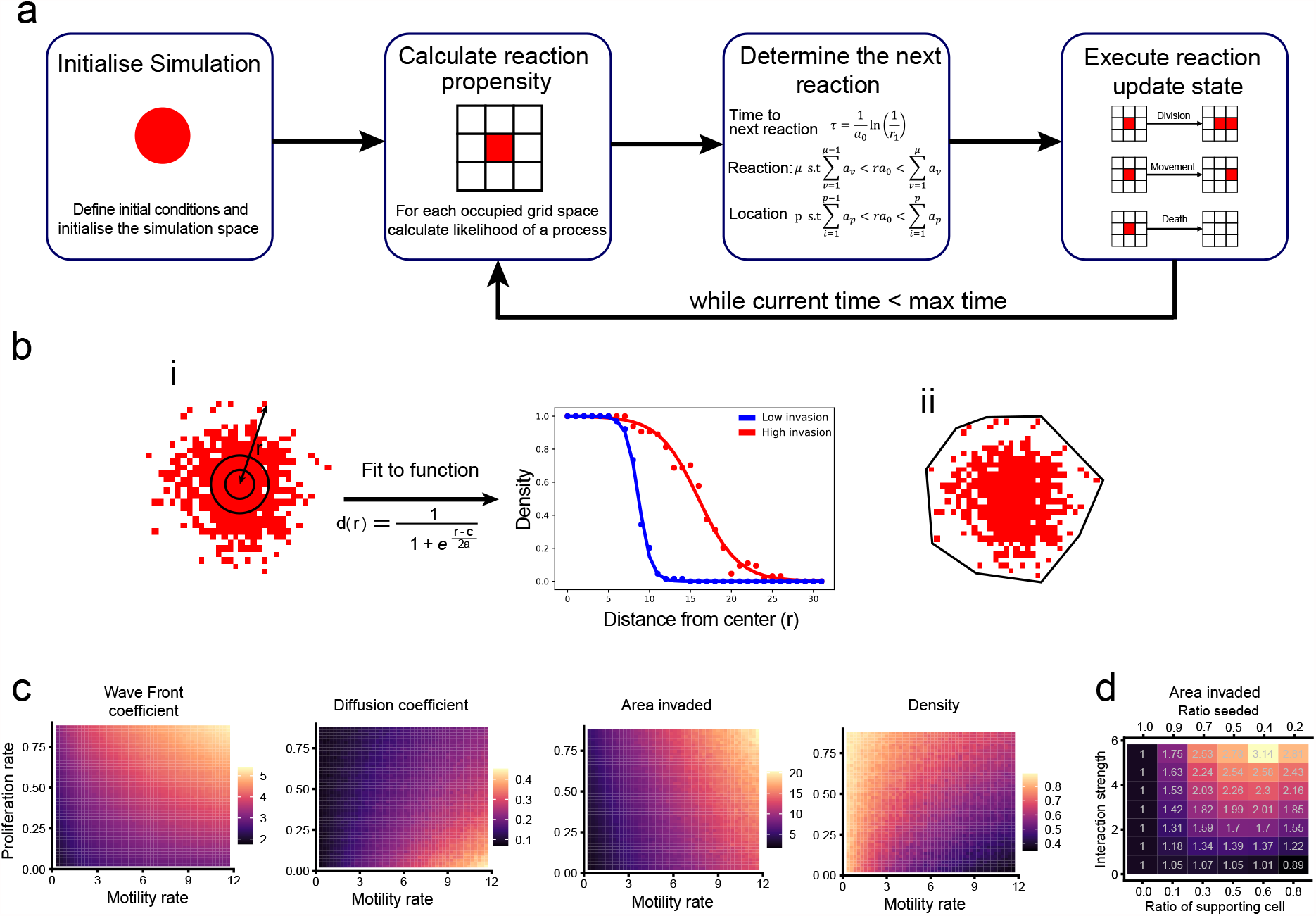
Simulating invasion. **a**, Illustration of simulation flow. **b**, Summary statistics used to measure invasion; i) Travelling wave solution to reaction diffusion equations fit to the spatial configuration of cells, ii) Convex hull area. **c**, Summary statistic of measurements of pure population invasion. Wave front coefficient, average distance invaded and area invaded have near orthogonal contours to density and diffusion coefficient. **d**, Highlighting the effect of introducing interactions on the phenotype observed. As the interaction strength increases, the value of the summary statistic measured (area invaded) also increase for the co-culture conditions. This demonstrates the benefit the interacting cell experience from being in a co-culture.

We use the travelling wave solution of a reaction-diffusion equation, commonly used to simulation glioma invasion^25^, in order to obtain diffusion and wave front coefficients (Figure 2bi). Using area invaded and average distance invaded as measures we are able summarise the extent of invasion, whilst density seeks to provided information on the spatial features of the area invaded (Figure 2bii, 2biii).

For a culture of a pure population with a fixed proliferation rate, the effect of increasing the motility rate is to increase the wave front coefficient, diffusion coefficient and area invaded whilst the density decreases (Figure 2c). This is not surprising as a population moving faster to occupy a larger area, whilst still growing at the same rate, will occupy larger area with the same number of total cells so this is less dense. On the other hand, keeping the motility rate fixed to monitor the effect increasing the proliferation rate, the wave front coefficient, area invaded and density all increase whilst the diffusion coefficient decreases (Figure 2c). A large number of cells possessing the same motility rate are indeed expected to occupy a larger area and more densely due to the increased growth rate.

Most importantly, the density and diffusion coefficient measurements have a near orthogonal contour to those of the wave front coefficient and area invaded. Thus, a pair of measurements from each group will help recover both rates (Figure 2c).

Finally, looking at a model with interactions we are able to see the effect of the seeding ratio and the interaction strength. In co-culture conditions as the interaction strength is increased, the summary statistic value also increases and therefore the invasive phenotype is enhanced. This is not reflected in the case of pure populations as there are no interclonal interactions (Figure 2d).

### Measuring the parameters that govern invasion of a ‘monoclonal’ population

In order to assess interactions between distinct subpopulations that leads to a change in phenotype, it is essential to understand the phenotype of a population in isolation.

Between the two clones isolated from the tumour SU-DIPG-VI, VI-E6 and VI-D10, VI-D10 appears to display a much more invasive phenotype as demonstrated by the summary statistics over time (Figure 3a) and imaging (Figure 3b). For example, both the diffusion and wave front coefficients increase faster for VI-D10 and thus using a travelling wave solution of a reaction-diffusion equation analysis leading to the conclusion that VI-D10 displays more prominent invasion than VI-E6. Similarly, between the two clones isolated from the tumour HSJD-DIPG-007, 007-F8 and 007-F10, 007-F10 displays a more invasive phenotype initially, as demonstrated by the summary statistics over time (Figure 3a) and imaging (Figure 3b). We do however, observe a plateau in the invasion of 007-F10 after day 2. These measures do not provide a single static parameter value, thus making quantitative comparisons difficult.

**Figure 3:**
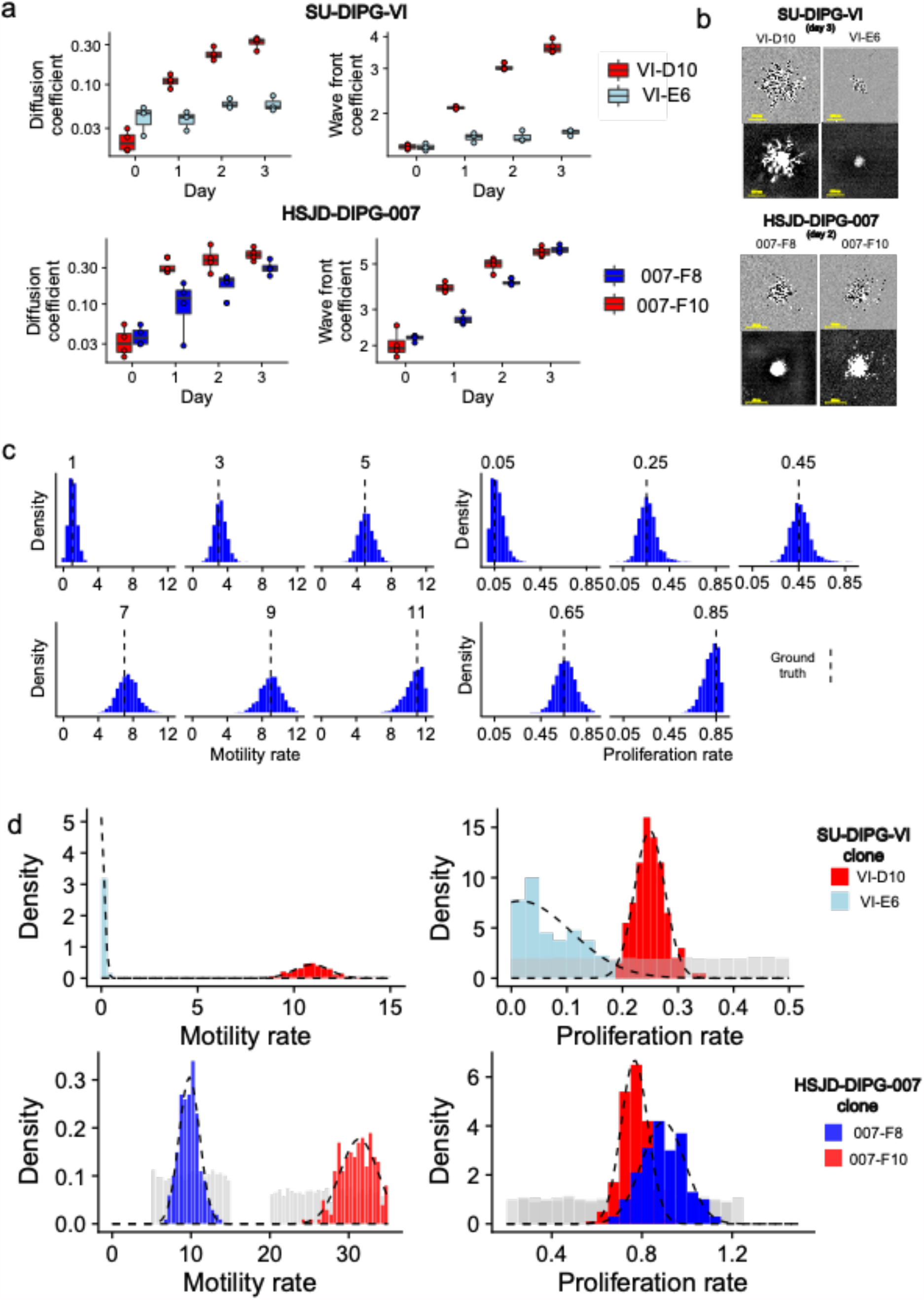
Measuring the parameters that govern invasion of a ‘monoclonal’ population. **a**, Summary statistics calculated for experimental images from SU-DIPG-VI clones VI-E6 and VI-D10 and HSJD-DIPG-007 clones 007-F8 and 007-D10, demonstrates a clear difference in the invasive phenotype. **b**, Images to highlight the differences in the growth of spheroid. **c**, Posterior distribution of simulated recovery of sample data with ground truth highlighted by a dashed vertical line. For a wide range of proliferation and motility rates the ground truth is recovered. **d**, Posterior distribution of parameters from experimental images, with truncated normal distributions fitted (dashed line). The prior distribution (grey) represents the distribution from which all simulation parameters were drawn from.

Using ABC inference, we first demonstrate that our scheme of summary statistics is able to accurately recover the invasion parameters from simulations. This is then followed by an application of the summary statistics to the experimental data, generating posterior distributions for our prediction of the motility and proliferation rates.

In order to validate the accuracy of our measurements in recovering invasion parameters, a set of sample simulations are run with pre-defined motility and proliferation rates. Summary statistics are then calculated for the sample simulations as well as each realisation in a simulation set of a large number of simulations with parameters drawn from random uniform distributions. The weighted Euclidean distance between the sample and simulation set is calculated for a single or combination of our summary statistics. The specific weights are explored using a genetic algorithm to find the most consistent recovery of the ground truth. Using a weighted measure of the difference between the wave front and diffusion coefficients observed in our sample simulations and each realisation, a distance is calculated between samples and the simulated databank. Simulations that have the closest summary statistics value to the sample data will be have their parameter values recorded and saved. From these simulations that we have saved, we are able to construct a posterior distribution of our parameter estimates. This posterior distribution represents the potential values we believe the parameter can take. Performing this analysis across a range of sample motility and proliferation rates, there is consistent recovery of the ground truth using this regime across a variety of proliferation and motility rates (Figure 3c). We also tested using an alternative combination of summary statistics, area invaded with density, and the recovery of parameter values is similar however with wider posterior distributions (Supplementary figure 1).

Applying this methodology to the binary masks generated from our invasion assays, the posterior distributions for the proliferation and motility rates we recover for SU-DIPG-VI clones VI-E6 and VI-D10 are consistent with the observation that VI-D10 displays significantly more pronounced invasion than VI-E6. The posterior distribution of proliferation and motility rates are then fitted to a distribution (truncated normal distributions), this allows for the quantification of the phenotype of each of our clones, as well as the ability to pass these distributions as prior assumptions for our mixed culture models (Figure 3d, Supplementary figure 1). An approach such as this allows for the propagation of uncertainty which often is lost when inferring a single value. This is crucial in avoiding skewing further inference.

We have also characterised the invasion parameters for the clones 007-F8 and 007-F10, from the bulk tumour HSJD-DIPG-007, showing that 007-F8 displays a similar rate of proliferation however a substantially reduced motile phenotype when compared to 007-F10 (Figure 3d).

### Measuring cellular interactions between distinct clones with differential invasion

Using the posterior distributions recovered for monoclonal populations, clones VI-E6 and VI-D10 from SU-DIPG-VI and clones 007-F8 and 007-F10 from HSJD-DIPG-007, we are able to condition our mixed culture simulations. Assigning motility and proliferation rates that are drawn from our posterior distributions, we can then focus on the effect that the interaction parameter has on the phenotype observed.

Highlighting a few simulations from a range of different interaction strengths, from low to high, we are able to understand how an interaction affects the invasive phenotype. In the absence of interactions, we can see that the normalised area decreases as the ratio the population is seeded decreases (Figure 4a). This is intuitive, as there have fewer cells to grow and move thus the invasion is less pronounced. However, introducing interactions this negative trend shifts to resembles a parabola with a peak that is increasing as the interaction strength increases. This suggests that introducing a small proportion of a supporting cell increases the invasive phenotype through interactions. However, it is important to remember that the cells also experience competition with one another for both space as well as nutrients, such as growth factors in the cell culture media, thus increasing the proportion of the supporting cell by too much leads to a plateau in the area invaded followed by a decline as competition increases. Competition begins to negate the effect of interactions with the benefit received achieving a plateau (Figure 4a).

**Figure 4:**
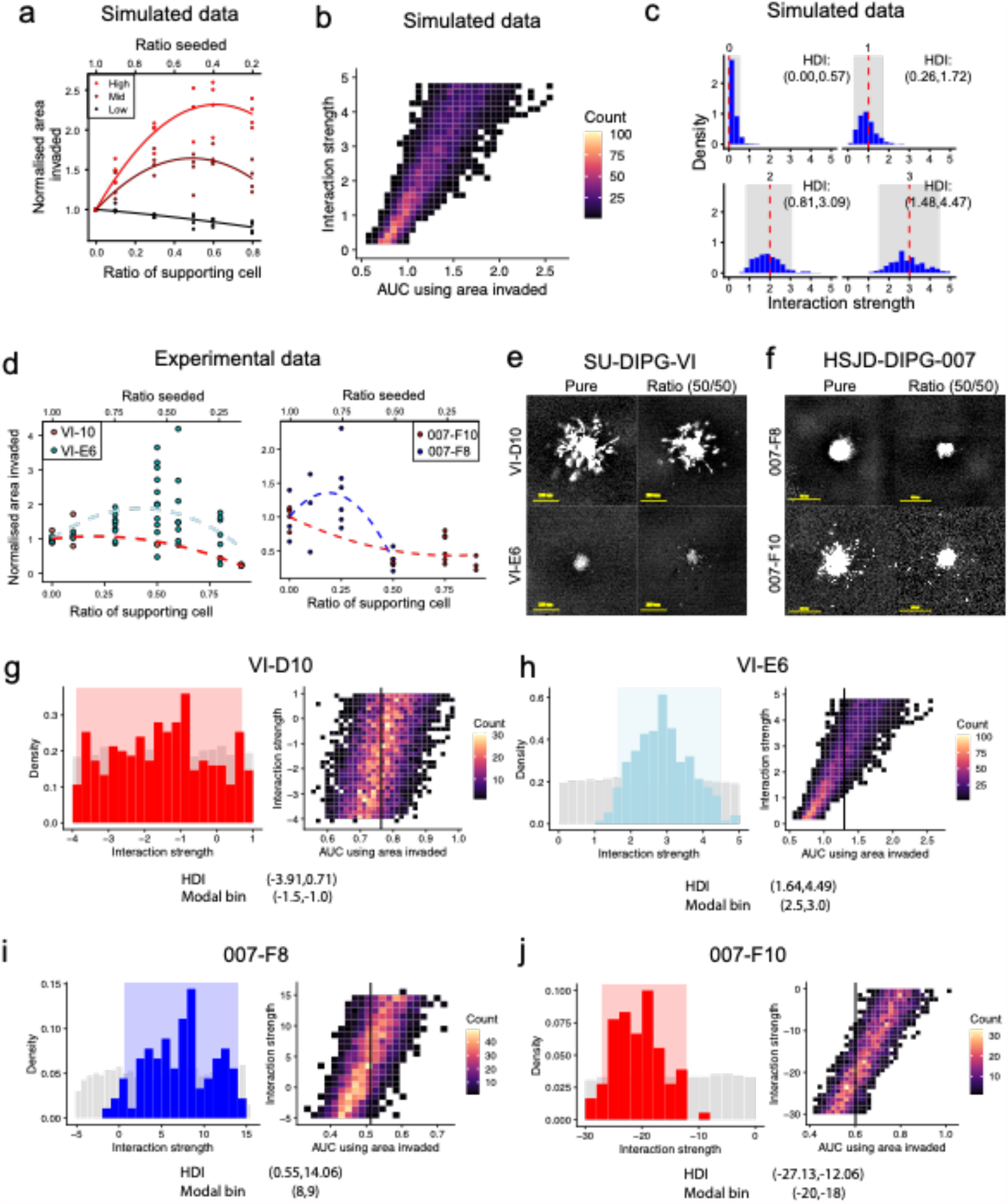
Measure cellular interaction between distinct clones with differential invasion. **a**, Summary statistics on simulated data conditioned on pure population parameters at a variety of interaction strengths. **b**, Linear relationship between area under curve and interaction strength. **c**, Posterior distribution of simulated recover from sample simulations, with ground truth (red dashed line) and credible interval (grey). **d**, Images demonstrate that VI-D10 displays lower invasion in a ratio than in isolations, whilst VI-E6 displays enhanced invasion **e**, Summary statistics on experimental data from day 3 demonstrates the parabolic relationship for VI-E6 but a decrease for VI-D10. **f**, Summary statistics on experimental data from day 2 demonstrates the parabolic relationship for 007-F8 but a decrease for 007-F10. In 007-F8 we see a similar parabolic relationship, however a decrease to the level of 007-F10 in the 0.5 ratio, while 007-F10 is strictly decreasing with a plateau. **g**, Posterior distribution for the interaction strength of VI-D10 (left) and simulated data plotted with vertical line representing experimental AUC (right). Showing interaction strength not credibly different from 0. **h**, Posterior distribution for the interaction strength of VI-E6 (left) and simulated data plotted with vertical line representing experimental AUC (right). Showing interaction strength credibly different from 0. **i**, Posterior distribution for the interaction strength of 007-F8 (left) and simulated data plotted with vertical line representing experimental AUC (right). Showing interaction strength credibly different from 0. **j**, Posterior distribution for the interaction strength of 007-F10 (left) and simulated data plotted with vertical line representing experimental AUC (right). Showing interaction strength credibly different from 0.

Noting that, between the different strengths of interactions, the curve fitting the summary statistic value vs ratio appears to become parabolic and shifting upwards. This can thus be summarised as an increase in the area under the curve, we can see that as the interaction strength increases the area under the curve (AUC) also increases (Figure 4b). This relationship should allow for the recovery of the interaction strength by using AUC as a summary statistic, obtained by using an approximate Bayesian computation methodology to generate posterior distributions.

Focusing on the effect on an interaction in a simulation, across the values of interaction strength sampled, there is a consistent recovery of the interaction strength parameter value. More importantly, we are able to confirm an interaction strength of zero, providing power to distinguish between the presence and absence of interactions. The ground truth is always contained in the credible interval for all samples and thus there is confidence in the ability to recover the interaction strength using AUC as a summary statistic (Figure 4c).

We can see in the experimental data, that the VI-D10 population displays results that look similar to that of no interactions whilst E6 to displays a parabolic phenotype which is more akin to the presence of an interaction (Figure 4d). Indeed, experimental images demonstrate that, VI-D10 invasion appears to be lower in the 50:50 ratio when compared to a pure culture. However, on closer inspection of VI-E6, there are more cells that are able to escape and invade the surrounding matrix in the 50:50 ratio when compared to the pure culture (Figure 4e). Similarly, we can see that between the clones 007-F8 and 007-F10, 007-F8 displays a weak parabolic relationship that quickly declines to below 1 at a ratio seeded of 50% whilst 007-F10 displays a significantly negative trend as the ratio seeded decreases (Figure 4f). Once again, closer inspection of images shows that indeed 007-F8 and 007-F10 have invaded a smaller distance at the 50% ratio (Figure 4g)

Applying these statistics to the experimental images, we then generated credible intervals for our experimental data. For VI-D10, it is clear that under both measurements the credible interval overlaps with 0, with a modal interaction strength interval of (−1.5, -1.0) (Figure 4g). This indicates that there is no evidence to suggest there is a positive interaction received by VI-D10. More interestingly, in the case of VI-E6 the credible interval for all replicates does not contain 0, with one replicate demonstrating modal interaction strength interval of (1, 1.5) (Figure 4f, Supplementary Figure 2). The evidence indicates that a model with interactions better explains the data than a model without, coupled with the reproducibility of the interaction strength across multiple experiments strongly indicated that VI-E6 receives a commensal interaction from VI-D10.

Further investigating the clones 007-F8 and 007-F10, we see that 007-F8 displays a positive interaction strength with HDI that does not overlap with 0 and a modal interaction strength interval of (8, 9) whilst F10 displays a credibly negative interaction strength with a modal interaction strength of (−20, -18). The evidence indicates that a model with negative interactions better explains the data than a model without, this indicates that 007-F8 displays an exploitative interaction on 007-F10.

We can also take a look at the distribution of the motility rate plus interaction strength against just the motility rate on its own, to understand the maximal effect interactions can have on the motility of a clone. We observe that the VI-E6 demonstrates a noticeable positive shift in the distribution whilst VI-D10 has largely overlapping distributions with a slight negative shift (Supplementary Figure 3). The same conclusions can be drawn for 007-F8 and 007-F10, with 007-F10 showing a noticeable negative shift in the distribution whilst 007-F8 has largely overlapping distributions with a slight positive shift (Supplementary Figure 3).

We see larger stochasticity our experimental data compared to simulations, this is particularly noticeable for measurements of VI-E6, the clone receiving the interaction VI-E6. This can be explained by the interplay between interactions and competition, where there is the potential for spatial competition to mask the effect of any interaction. This occurs, when VI-D10 is able to quickly engulf VI-E6, thus limiting VI-E6 in terms of finding space to invade. We demonstrated this by overlaying the green fluorescent channel on top of the phase images, to highlight the possible scenarios, from complete freedom to invade in all directions to complete spatial restriction from being engulfed (Supplementary figure 4).

Here we have demonstrated that, from two pairs of subclonal populations derived from two different bulk tumours, the presence of a commensal interaction where one population is benefitting from being culture with the other as well as an exploitative interaction where one population benefits and another suffers from being cultured together.

## Discussion

In this study, we present a new quantitative methodology enabling the detection of positive spatial interactions that affect the invasive phenotype and quantify them. We applied this to a unique set of single cell derived clones from autopsy samples. These findings help understanding intra-tumoural heterogeneity, a phenomenon linked to the adverse prognosis for patients particularly in the case of paediatric high-grade gliomas^5^. The ability to detect and measure subclonal interactions can open the avenue for treatments that seek to contain tumours by disrupting positive interactions and promoting negative interactions.

Spatial interactions, are a largely unexplored in cancer, in particular those affecting cellular invasion, with much of the literature focusing on the proliferative interactions. Interactions in cancer are rarely identified, as such it would be interesting to see this work recreated in a larger number of samples. Whilst we have demonstrated the ability to detect positive and negative interactions as well as estimate their strengths across biological replicates, there is significant room for further developments. Detecting interactions is merely the initial step, the ultimate goal should be to use our understanding of such interactions to improve patient outcomes such as through therapeutic interventions. This study combines computational modelling study of intra-tumoral interactions between subclones and created a model trained from two distinct patient-derived cell lines well characterized phenotypically and genotypically. This new model can therefore be applied to more cell lines and subclones, and potentially more cancer types. Whilst, also focusing on determining mechanisms that underpin such interactions will enable for the development of therapeutic targets that leverage interclonal interactions to improve patient prognosis.

In the case of this study, a key extension would be to introduce mechanistic modelling in order to determine the nature of interactions and find the causal factor. There are a multitude of different avenues for interactions to exhibit themselves, such as being pulled via cell-cell adhesion or chemotaxis across a chemical gradient. This will require for the creation of assays tailored to detecting these interactions which should be matched with computation models that aim to do the same. Models to recreate the interplay between spatial competition and interactions will help broaden the understanding of how likely a particular clone is to be engulfed and how this changes with the ratio seeded and strength of interaction.

Whilst the model used in this study is relatively simple, as it does not include the presence of resource concentration gradients, secreted factors as well as a multitude of other considerations, it provides a crucial stepping stone in detecting interactions. Since positive interactions are rare, our approach allows for the identification of interactions which can in turn be an indication of where resources and time should be spent to further explore the causes of such interactions.

## Materials and methods

### Cell culture

Patient-derived cultures SU_DIPG-VI (*H3F3A* ^K27M^, *TP53*^p.R175H & p.E198*^, *MYC* ^amp^) and HSJD-DIPG-007 (*H3F3A* ^K27M^, *ACVR1* ^R206H^, *PPM1D*^p.P428fs▲^, *PIK3CA*^p.H1047R^) were grown in stem cell media consisting of Dulbecco’s Modified Eagles Medium: Nutrient Mixture F12 (DMEM/F12), Neurobasal-A Medium, HEPES Buffer Solution 1M, sodium pyruvate solution 100nM, nonessential amino acids solution 10mM, Glutamax-I Supplement and penicillin Streptomycin solution (all Thermo Fisher, Loughborough, UK). The media was supplemented with B-27 Supplement Minus Vitamin A, (Thermo Fisher), 20ng/ml Human-EGF, 20ng/ml Human-FGF-basic-154, 20ng/ml Human-PDGF-AA, 20ng/ml Human-PDGF-BB (all Shenandoah Biotech, Warwick, PA, USA) and 2µg/ml Heparin Solution (0.2%, Stem Cell Technologies, Cambridge, UK) to constitute the complete media. In brief, cells were incubated at 37°C, 5% CO2, 95% humidity and were refed at least twice weekly with complete media. Cell authenticity was verified using short tandem repeat (STR) DNA fingerprinting. Further details on human glioma lines used can be found on www.crukchildrensbraintumourcentre.org/research/resources/cell-line-repository/.

### Cell labelling

Different clones were derived from the bulk of SU-DIPG-VI (VI-D10 and VI-E6) and HSJD-DIPG-007 (007-F8, 007-F10) as previously described in our previous publication^5^.

Each clone was stably labelled with the lentiviral “gene ontology” (LeGO) vectors^26^. Transduction were performed using lentivirus encapsulated with the following plasmids: the plasmid Lego-iC2, mCherry expressing vector (#27345; Addgene) or the plasmid Lego-V2, Venus expressing vector (#27340; Addgene).

Briefly, each LeGo plasmid was transfected into HEK293T cells together with the Trans-Lentiviral shRNA Packaging System (#TLP5912; Dharmacon) helped by Lipofectamine 2000 (#11668030; Invitrogen). Forty-eight hours post-transfection viral particles in the supernatant were collected, filtered through Millex-HV 0.45um filter (#SLHVM33RS, Millipore), concentrated with LentiX Concentrator according to the manufacturer’s instructions (#631231, Takara) then stored aliquoted at -80C. For a clonal selection, the transfected cells were single cell flow sorted into the inner 60 wells of 96 well plates ultra-low attachment round bottom (#7007, Corning) using a Beckman Coulter MoFlo in Class IIA2 biohazard containment hood. Cells were dropped in 100µl/well of complete media supplemented with 2X growth factors, Primocin (#ant-pm, InvivoGen), Plasmocin (#ant-mpt, InvivoGen), penicillin and streptomycin (Life Technologies). In order to enhance the fluorescent signal, clones were also labelled transiently with NucLight Red or Green BacMam 3.0 Reagent (#4621 and 4622 respectively; Sartorius (discontinued) before seeding for the assays.

### Migration/invasion assays

50-100μm diameter neurospheres were harvested after three days culture of the SU-DIPG-VI and HSJD-DIPG-007 generated clones and were used for the invasion and migration assays. Invasion assays were performed as previously described ^5,27,28^, with some modifications. Briefly, a total of 100μl medium was removed from ULA 96-well round-bottomed plates containing neurospheres of 100μm in diameter. For the invasion study, 100μl of matrigel was gently added to each well (6 replicates) and plates were incubated at 37°C, 5% CO2, 95% humidity for 1hr. Once the matrigel solidified, 100μl/well of culture medium was added on top.

3D migration assays were similarly performed as previously described^5,27,29^, with some modifications. Briefly, flat-bottomed 96-well plates (Greiner Bio-one) were coated for 2hrs at room temperature with 50μl/well of fibronectin, laminin (Sigma-Aldrich) 10μg/ml in PBS with calcium and magnesium, or 125μg/ml matrigel (Corning) in culture medium in absence of growth factors. Once coating was completed, a total of 200μl/well of culture medium was added to each well. A total of 100μl medium was removed from ULA 96-well round-bottomed plates containing neurospheres of 50-100μm in diameter, and the remaining medium including the neurosphere were transferred into the pre-coated plates. Starting from time zero, and at 3h intervals up to 6 days, automated image analysis was carried out on the live-cell imaging system IncucyteS3 using the *Whole Well* application.

### Imaging

Images are taken using the Incucyte S3 live-cell analysis system using the spheroid scanning module with a 4x objective. Images were taken of the phase red, and green imaging channels.

### Image segmentation

In first step, images were exported with out of focus images discarded and pre-processing consisting of CLAHE (contrast-limited adapted histogram equalisation) applied. This was applied on a 5×5 pixel window, resulting in enhancement of the signal in the fluorescent channels.

We then prepared training data, in order to train a neural network to identify features from the images to segment on. This was a labour-intensive process, in which we create ground truth training masks for learning. These masks were created for both the phase and green fluorescence channel. We used the training data to train a ResNet-UNet neural network, using an 80-20 training-validation split and running the training process until the validation error converged. This trained model was used for the segmentation in the next step.

We then applied our segmentation algorithm on the phase and green fluorescent images resulting in raw mask that represented a score on how likely a pixel contains a cell. We used a cut off of 0.5, above which we deemed a pixel to be positive and negative below. This converted the raw masks into binary marks.

Following this we removed the effect of slight fluctuations in the position of the microscope lens, which resulted in images that may be slightly shifted. In order to do this, we used MATLAB to perform image registration. This is performed using the phase channel to generate a transform which shifts an image in the x-y plane to fit the previously seen image. Sequentially in order of time, we applied this transform to all images and all channels resulting in images that were overlaid perfectly on one another. Now that all images are registered we used the phase masks to filter any background fluorescence we may have picked up in the green channel. This approach essentially checks if a spot is both phase and green positive in order for it to be included in analysis.

We then reviewed each binary mask (by creating an outline) to review their accuracy, as there is a difference in performance of this algorithm depending on the size of cells and intensity of label expression. Our segmentation approach was overly cautious and as such we have in some conditions a significant number of false negative pixels. We corrected for these by comparing images with the mask. Following this, any inaccurate masks are corrected for and a final binary mask is created.

### Genetic algorithm

These are a useful class of algorithm for parameter selection. The approach involved creating a fitness function on which parameter values will be compared against. We sought to find the parameter values with the maximum fitness values. This involved initialising simulations with many different parameter values and based on fitness only allowing a subset of these to be carried forward. These carried forward values are perturbed to explore locally and find fitter parameter values. Successive iterations of this led to a convergence in the mean fitness of accepted parameter values and we extracted the maximum fitness parameter value from this. In order to limit fitting to local optima we also generated an estimated trajectory of the fitness function and check to see our parameter aligned with the global minima. Analysis was produced using the GA package in R.

### Bayesian computation

We used an approximate Bayesian computation approach throughout this study. The general approach here was to create a large databank of simulations. Each realisation in our databank is initiated from a set of parameters, where our parameters of interested was drawn from a non-informative uniform random distribution. In order to reduce the size of the databank, an exploratory simulation was run to validate the limits of the prior to ensure it encompasses the posterior. For a sample, either experimental or simulated measurement, we calculated the distance between this and each realisation in our databank using a weighted Euclidean distance (derived using a genetic algorithm). We set an acceptance threshold, any distances that fell below this value were accepted and were used to generate a posterior distribution.

In theory, for an infinitely large databank, as the threshold approaches zero the posterior distribution converges to a single value, this value is the true parameter value.

For mono-culture simulations we fit a truncated normal distribution to the posterior distribution using the packages fitdistrplus, truncnorm and extraDistr in R.

However, in practice this is computationally infeasible and as such we generated credible intervals. These are the 95% HDI of the posterior distribution, and we expect our true parameter value to lie in this interval, this calculated using the HDInterval package in R.

## Supplementary information

### Agent based cellular automaton

Cellular invasion is the process whereby cells penetrate into their neighbourhood, this is driven by the proliferation rate of cells as well as their motility. Previous models of cellular invasion are able to encompass both factors, an example of one such model is the use of reaction-diffusion equation models. These models have been used extensively to model glioma invasion and we can see from the travelling wave solutions^25^, that the invasive phenotype is determined both by the motility and proliferation rates. These models have the advantage of being computational inexpensive however, do not lend themselves to be extended to more complex models such as those encompassing multiple subpopulations or with significant spatial considerations, as this often leads to systems with inconsistent behaviour. As such, we utilise cellular automata (CA) mimic the invasion of cells in-silico to capture the spatial restrictions and interactions we seek to model between distinct subpopulations.

A cellular automaton is a class of discrete models in computing, consisting of a finite-dimensional grid which each point in the grid representing a finite state and pre-determined rules that are followed. These models have been used extensively to study natural processing including models in oncology^30^.

We created a CA model on a three-dimensional grid with each point representing an empty space or containing a cell. Each cell can undergo a pre-determined set of possibilities; divide into a neighbour grid point, move to a neighbouring grid point or undergo cell death (Figure 2A). Each of these processes occur at different rates and as the model is extended we will have a distinct set of rates for each individual subpopulation we are modelling. To determine the next process to occur at any given time and the location at which this occurs we employed Gillespie’s stochastic simulations algorithm (SSA)^24^. This is a common choice^30^ and has been shown to generate a statically consistent trajectory of stochastic equations, with a caveat of being computationally intensive.

We initiated a simulation with cells set to a pre-determined radius around a central point. From this state, we calculated the propensity for each process to occur. Next, we chose a random process and cell for the next process to occur, weighted for propensity, as well as the time to next reaction using the total propensity. At this point we updated the state of the simulation by updating the current time and executing the process selected. This loop is repeated until the end time of the simulation is achieved, at set intervals we take a snapshot of the state of the simulation to use for analysis (Figure 2A).

We also seek to model the invasion of two distinct subpopulations cultured together. In this model, we have to accounted for the ratio of each subpopulations as well as any interactions present that affect the motility rate. A model such as this is crucial as it informs on what we expect to see in the absence of interactions. This is difficult to determine solely from in vitro experiments as we are unable to rule out the presence of interactions and have no basis to guide our observations. We can set the interactions between the two subpopulations to be zero, so neither population affects the rates of the other. This will serve as our null model of no interactions.

We modify our previous model to by allowing for the presence of three different states in our simulations; empty, cell of type A or cell of type B. The rest of the simulation is carried out in the same manner as previously described. We now have an additional parameter which determines the initial ratio between the two subpopulations. We introduced an interaction strength parameter which affects the motility rate of a cell and it is scaled according to the proportion of another cell. The interaction strength is now defined by; Mi = rj * Iij where subscripts dictate the cell, M is the final motility rate of a cell, r is the current proportion and Iij is the interaction strength on i received by j. We have chosen this regime as it is a simple method of introducing interactions and rj can be scaled according to the distance of interaction, which currently we keep unlimited. These longer-range interactions can be more akin to chemokine signalling as opposed to shorter range contact induced interactions. Whilst there are a multitude of different regimes to implement, we are drawing our attention to distinguishing that a model with no interactions is insufficient in explaining the data.

These simulations were written in CUDA C.

---

**Supplementary figure 1:**
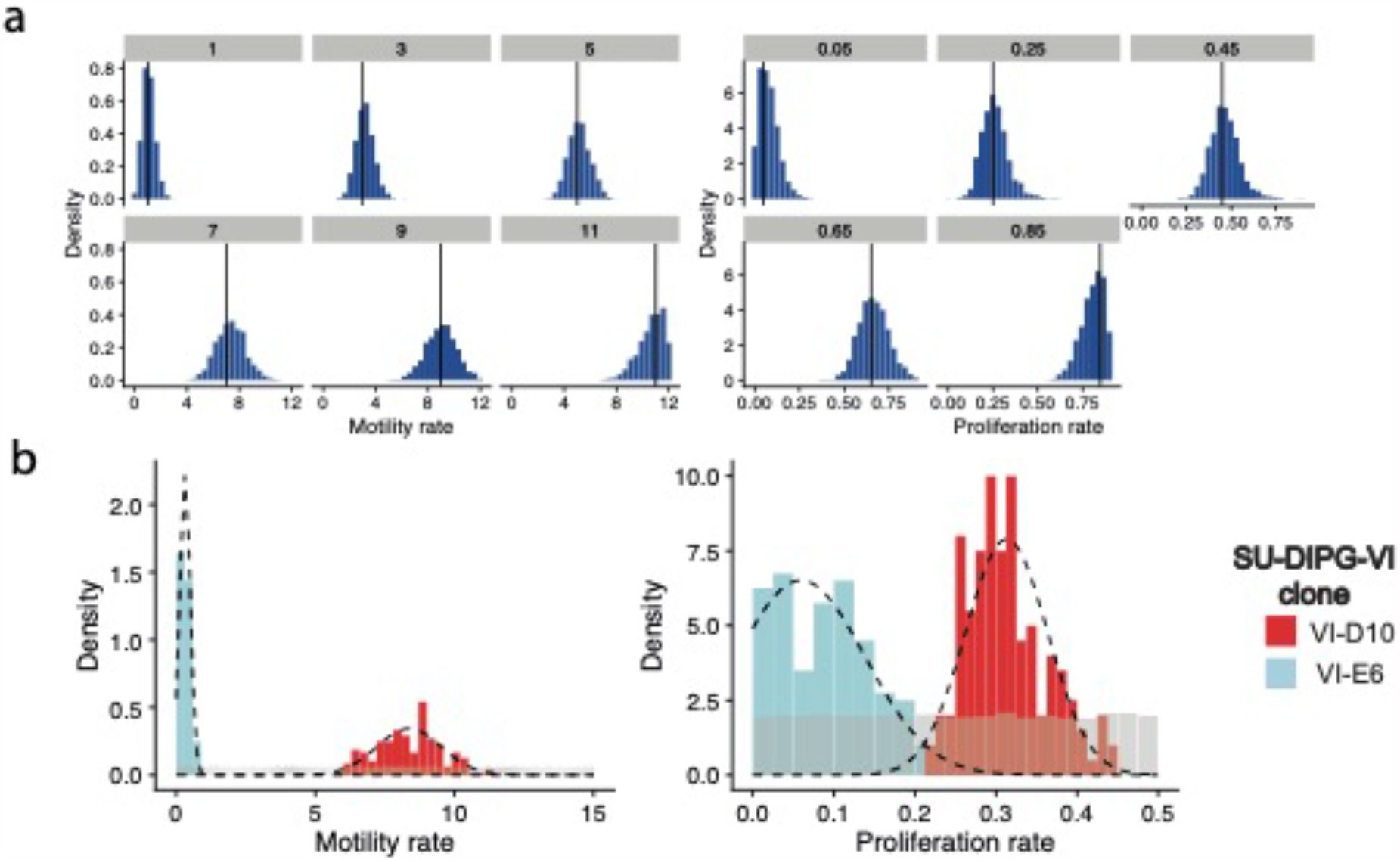
Recovery of invasion rates for SU-DIPG-VI clones VI-E6 and VI-D10 using area invaded with density of invaded area as summary statistics. **a**, Simulated recovery. **b**, Experimental recovery.

**Supplementary figure 2:**
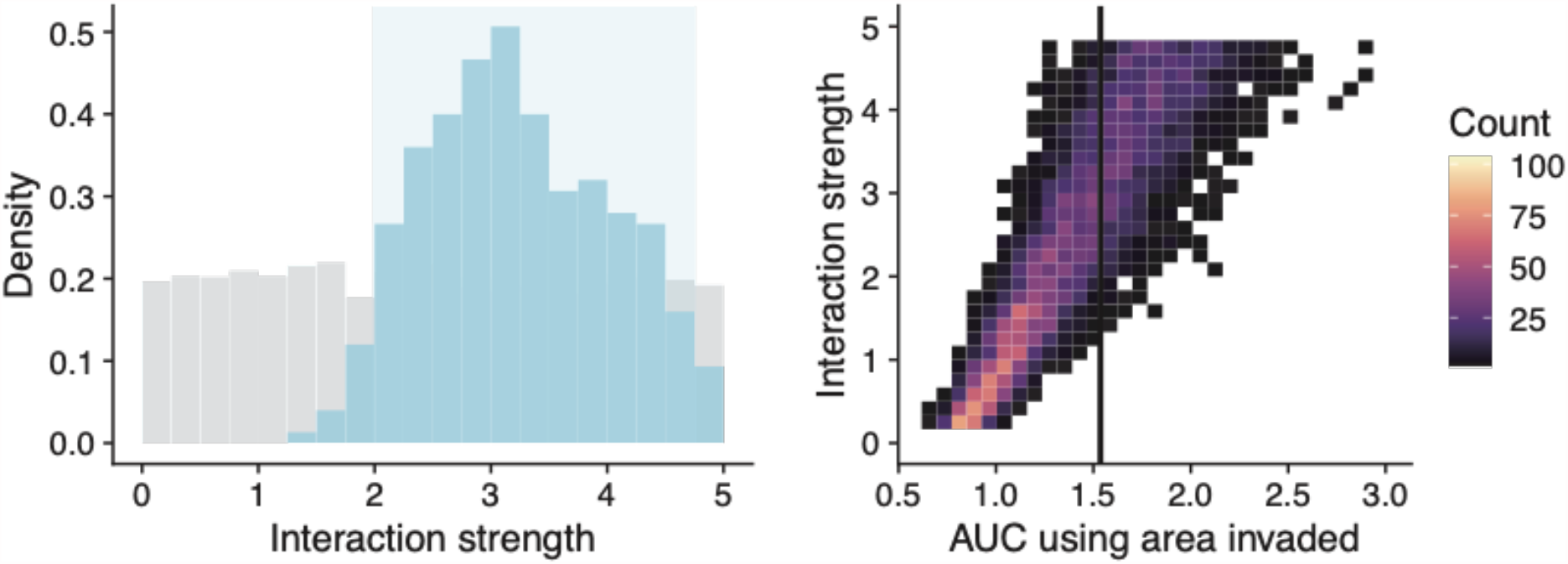
Additional replicate validates the experimental recovery of a positive interaction observed for SU-DIPG-VI clone VI-E6.

**Supplementary figure 3:**
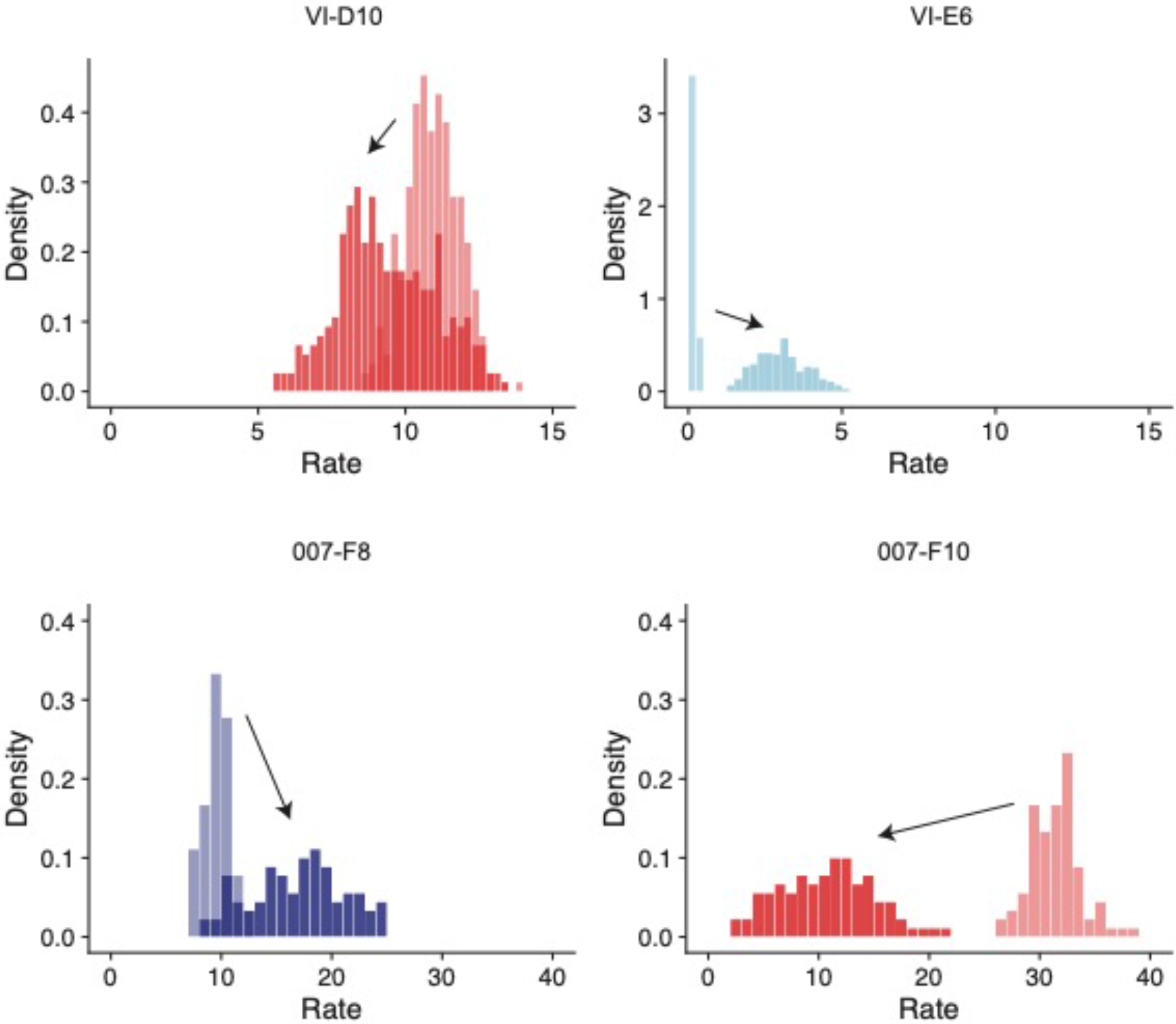
Distribution of motility rate and motility rate plus interaction strength highlights the effect of an interaction on the motility rate of a clone.

**Supplementary figure 4:**
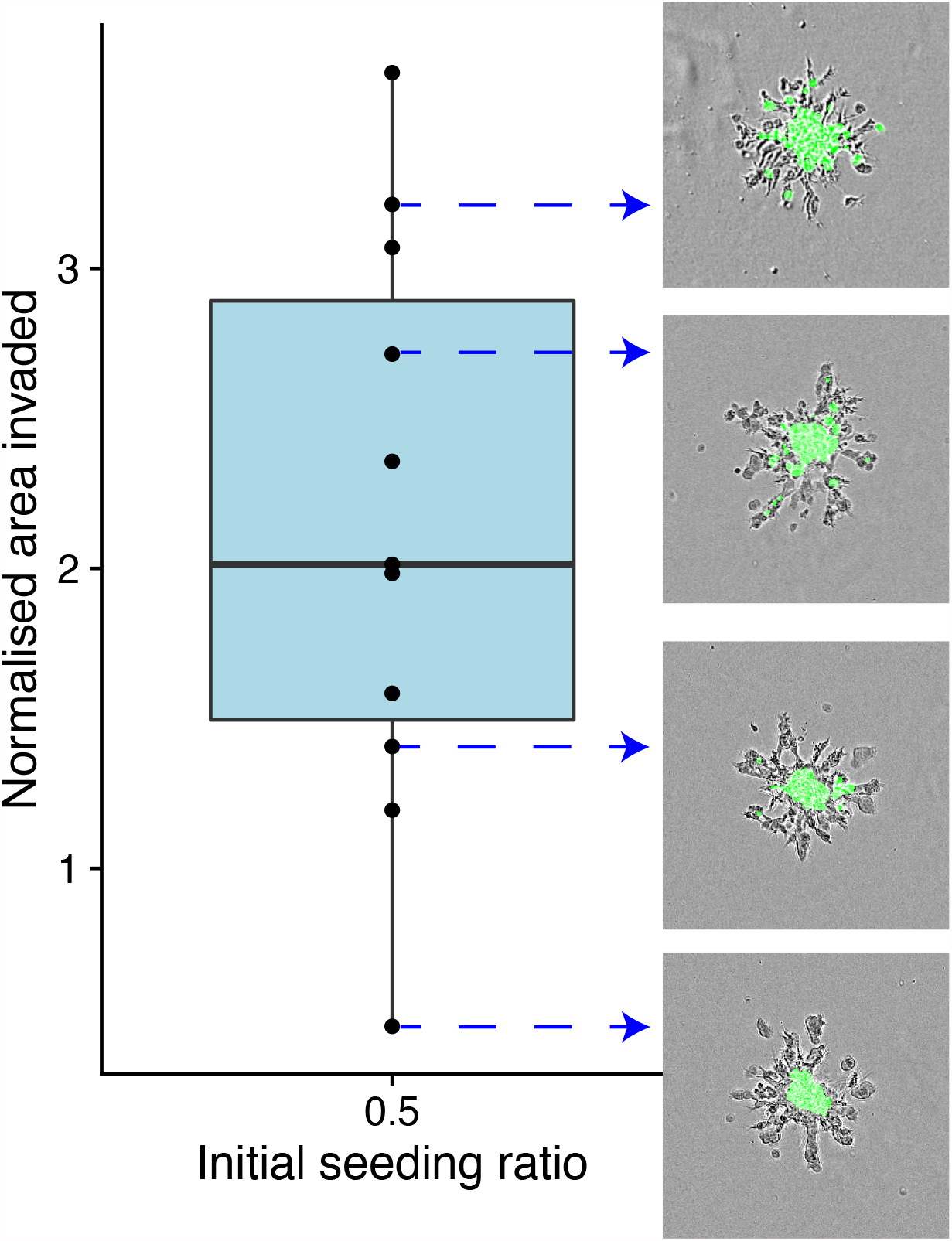
The interplay between spatial competition and positive interactions leads to variable outcomes in the invasive phenotype observed. Using SU-DIPG-VI E6 as an illustrative example.

